# The complete mitochondrial genome of a parthenogenetic ant *Monomorium triviale* (Hymenoptera: Formicidae)

**DOI:** 10.1101/2021.03.02.433554

**Authors:** Naoto Idogawa, Chih-Chi Lee, Chin-Cheng Scotty Yang, Shigeto Dobata

**Affiliations:** Laboratory of Insect Ecology, Graduate School of Agriculture, Kyoto University, Kyoto 606-8502, Japan; Institute of Evolution, Department of Evolutionary and Environmental Biology, University of Haifa, Haifa, 31905, Israel; Department of Entomology, Virginia Polytechnic Institute and State University, Blacksburg, VA, 24061-0131, USA; Department of General Systems Studies, Graduate School of Arts and Sciences, The University of Tokyo, Komaba, Meguro, Tokyo 153-8902, Japan

**Keywords:** Hymenoptera, Formicidae, thelytoky, *Monomorium*, mitogenome

## Abstract

*Monomorium* is one of the most species-rich yet taxonomically problematic ant genus. An East Asian species, *M. triviale* Wheeler, W.M., 1906, are reproduced by obligate thelytokous parthenogenesis and performs strict reproductive division of labor. We sequenced the *M. triviale* mitogenome using next-generation sequencing methods. The circular mitogenome of *M. triviale* was 16,290 bp in length, consisting of 13 protein-coding genes, two ribosomal RNA genes, 22 transfer RNAs, and a single non-coding region of 568 bp. The base composition was AT-biased (82%). Gene order rearrangements were detected and likely to be unique to the genus *Monomorium*. We announce the *M. triviale* mitogenome as additional genomic resources for elucidating phylogenetic and taxonomic problems of *Monomorium* and comparative genomics of parthenogenetic ant species.

---

In the hyperdiverse ant subfamily Myrmicinae, *Monomorium* Mayr, 1855 is one of the most species-rich genera with over 300 described species including several successful tramp species such as the flower ant *M. floricola* and *M. salomonis*, and the pharaoh ant *M. pharaonis* (Pontieri & Linksvayer 2019). However, recent studies (Ward 2015, Sparks 2019) suggest polyphyly of this genus, and genomic resources are therefore essential for resolving such a taxonomic issue.

An East Asian species, *M. triviale* Wheeler, W.M., 1906 is particularly of our interest as it reproduces by thelytokous parthenogenesis where virgin queens produce both workers and next-generation queens (Gotoh at al. 2012; Idogawa et al. 2021). To date, only partial mitochondrial DNA sequences have been reported, with all of which being identical among populations in Japan (Idogawa et al. 2021). Hence, a complete mitochondrial genome of this species can provide additional information for further analysis. Here, we present the first complete mitogenome for *M. triviale*.

A colony of *M. triviale* headed by a single queen was collected in Takaragaike Park, Kyoto, Japan (35.060087 N, 135.788488 E) on Sept. 9, 2017. The queen and her parthenogenetic offspring produced later in the laboratory (larvae and pupae, approx. 100 individuals) were fixed in 99.5% EtOH. We extracted genomic DNA from the pooled individuals with DNeasy Blood and Tissue kit (Qiagen, Hilden, Germany). We sequenced the pooled DNA on the HiSeq X sequencer (Illumina, San Diego, CA) at Macrogen Japan Corp., Tokyo. After removing adapters with Trimmomatic v0.39 (Bolger et al. 2014), we conducted de novo mitogenome assembly based on 4,327,159 paired-end sequence reads using NOVOPlasty v3.6 (Dierckxsens et al. 2017), with *Solenopsis invicta* mitogenome (NCBI accession: NC_014672) as a seed. Average read coverage of the mitogenome assembly was 37,840, providing ample depth for correctness. We annotated protein-coding genes (PCGs), rRNAs and tRNAs using GeSeq (Tillich et al. 2017), MITOS (Bernt et al. 2013), NCBI ORF-finder (Rombel et al. 2002), and ARWEN (Laslett and Canbäck 2008). The sequence information was deposited in the DNA Data Bank of Japan under accession number: LC605004. A specimen was deposited at the Laboratory of Insect Ecology, Graduate School of Agriculture, Kyoto University (http://www.insecteco.kais.kyoto-u.ac.jp; N. Idogawa; under the voucher number Mtri_20170909_4).

The complete mitogenome of *M. triviale* was 16,920 bp in length, which is comparable to those of other ant species. The nucleotide composition was AT-biased (82%). The mitogenome contained 13 protein-coding genes (PCGs), two rRNAs, and 22 tRNAs, which are typical for most metazoan animals. All PCGs used ATG or ATT as the start codon and TAA or TAG as the stop codon. The tRNAs, ranging in size from 59 to 74 bp, were similar to those of other ants (54 to 90 bp). The control region presumably corresponded to the single largest non-coding AT-rich region (569 bp, A+T 94%). Gene order of *M. triviale* was identical to that of a congener *M. pharaonis* (NC_051486.1). Notably, the gene order of the two *Monomorium* species had two gene rearrangements: an inversion between *trn*P and *ND1* (Myrmicinae, *ND6-CYTB-trn*S; *Monomorium*, <u>*trn*S</u>-<u>*CYTB*</u>-<u>*ND6*</u>; underlines indicate inverted genes) and translocations between *ND3* and *trn*F (Myrmicinae common, *trn*A-*trn*R-*trn*N-*trn*S-*trn*E; *Monomorium*, *trn*R-*trn*E-*trn*A-*trn*N-*trn*S). This feature was different from the common gene order of the subfamily Myrmicinae and likely unique to *Monomorium* ants (Babbucci et al. 2014; Park 2020). This may help identify the genus *Monomorium sensu stricto*, in addition to nucleotide substitutions.

We inferred the phylogenetic relationships of 25 ant species using the concatenated nucleotide sequences of all 13 PCGs, with the honeybee *Apis mellifera* as an outgroup. Sequence alignment was constructed using ClustalW (Thompson et al. 2003) implemented in MEGA-X (Kumar et al. 2018). The GTR + I + G model was determined as a best-fit model by ModelTest-NG v0.1.6 (Darriba et al. 2020). Both a maximum likelihood tree made by RAxML-NG v1.0.0 (Kozlov et al. 2019) and Bayesian inference tree made by MrBayes v3.2.7 (Ronquist et al. 2012) consistently support the current phylogenetic placement of *Monomorium* in the subfamily Myrmicinae (Figure 1).

**Figure 1.**
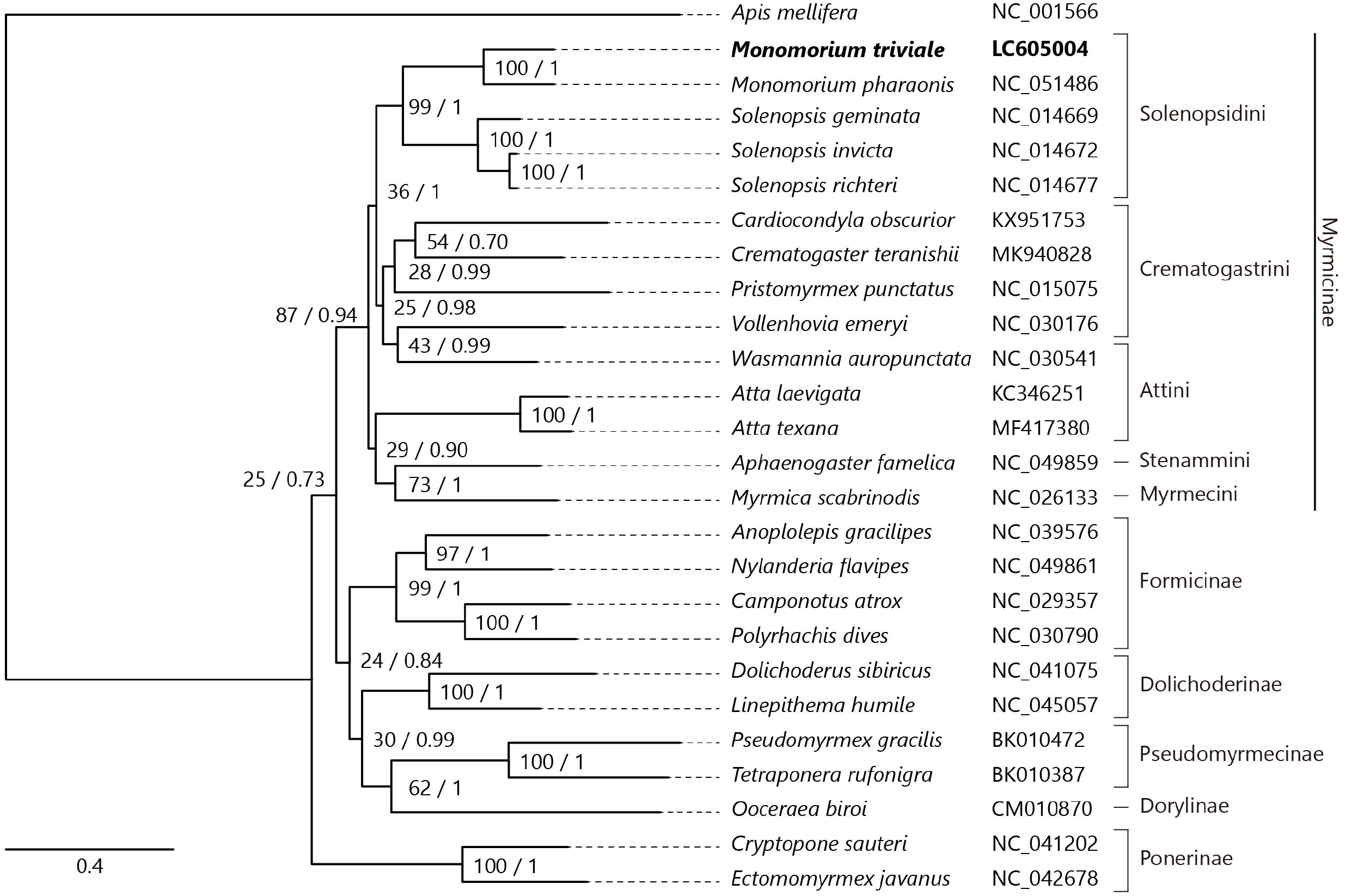
Maximum likelihood (1,000 bootstrap repeats) and Bayesian inference (100,000 generations) trees showing phylogenetic relationships among 25 ant species. A honeybee, *Apis mellifera* was used as an outgroup. Phylogenetic tree was drawn based on maximum likelihood tree. *Monomorium triviale* in bold is the result obtained in this study. The numbers above branches indicate bootstrap support values for maximum likelihood tree and posterior probability for Bayesian inference tree, in order.

In conclusion, the newly sequenced complete mitochondrial genome of *M. triviale* provides additional resources for further phylogenetic characterization of the taxonomically problematic genus *Monomorium* and comparative genomics of parthenogenetic ant species.

## Data availability statement

The genome sequence data that support the findings of this study are openly available in DDBJ/GenBank at (https://www.ddbj.nig.ac.jp) under the accession no. LC605004. The associated BioProject, SRA, and Bio-Sample numbers are PRJDB12079, DRX301164, and SAMD00394597 respectively.

## Acknowledgments

We are grateful to Kenji Matsuura who allowed us to use his laboratory.

